# Neglected no longer: Phylogenomic resolution of higher-level relationships in Solifugae

**DOI:** 10.1101/2022.10.22.513338

**Authors:** Siddharth S. Kulkarni, Hugh G. Steiner, Erika L. Garcia, Hernán Iuri, R. Ryan Jones, Jesús A. Ballesteros, Guilherme Gainett, Matthew R. Graham, Danilo Harms, Robin Lyle, Andrés A. Ojanguren-Affilastro, Carlos E. Santibañez-López, Gustavo Silva de Miranda, Paula E. Cushing, Efrat Gavish-Regev, Prashant P. Sharma

## Abstract

Considerable progress has been achieved in resolving higher-level relationships of Arthropoda in the past two decades, largely precipitated by advances in sequencing technology. Yet, dark branches persist in the arthropod tree of life, principally among groups that are difficult to collect, occur in cryptic habitats, or are characterized by minute body size. Among chelicerates, the mesodiverse order Solifugae (commonly called camel spiders or sun spiders) is one of the last orders of Arachnida that lacks a higher-level phylogeny altogether and has long been characterized as one of the “neglected cousins”, a lineage of arachnid orders that are comparatively poorly studied with respect to evolutionary relationships. Though renowned for their aggression, remarkable running speed, and adaptation to arid habitats, inferring solifuge relationships has been hindered by inaccessibility of diagnostic characters in most ontogenetic stages for morphological datasets, whereas molecular investigations to date have been limited to one of the 12 recognized families. In this study we generated a phylogenomic dataset via capture of ultraconserved elements (UCEs) and sampled all extant families. We recovered a well-resolved phylogeny of solifuge families, with two distinct groups of New World taxa nested within a broader Paleotropical radiation. To provide a temporal context to solifuge diversification, we estimated molecular divergence times using fossil calibrations within a least-squares framework. Solifugae were inferred to have radiated by the Permian, with divergences of most families dating to the post Paleogene-Cretaceous extinction. These results accord with a diversification history largely driven by vicariance as a result of continental breakup.

## Introduction

Chelicerata is an ancient, monophyletic group of arthropods that is characterized by extensive diversity, high body plan disparity among their different orders, and inclusion of numerous charismatic taxa, such as spiders, scorpions, and horseshoe crabs. Despite their considerable diversity, nearly every group of chelicerate orders has benefitted from recent advances in molecular phylogenetics, genomics, and renewed interest in evolutionary dynamics. The past decade alone has witnessed the first molecular phylogenetic hypotheses for several orders, such as Scorpiones, Ricinulei (hooded tick-spiders), Palpigradi (microwhip scorpions), Thelyphonida (vinegaroons), Schizomida (short-tailed whip scorpions), and Amblypygi (whip spiders) (Murienne et al. 2013a; Giribet et al. 2014; Fernández and Giribet 2015; Sharma et al. 2015; Clouse et al. 2017). More diverse chelicerate groups, such as Acari (mites), Araneae (spiders), Opiliones (harvestmen), and Pseudoscorpiones, have witnessed a surge of evolutionary inquiry in the past ten years (Hedin et al. 2012; Bond et al. 2014; Fernández et al. 2014; Garrison et al. 2016; Fernández et al. 2017; Klimov et al. 2017; Santibáñez-López et al. 2019; Benavides et al. 2019a; Kulkarni et al. 2020; Ballesteros et al. 2021; Kallal et al.2021; Santibáñez-López et al. 2022). This decade is also notable for the completion of the first genomes for several chelicerate orders (Sanggaard et al. 2014; Gulia-Nuss et al. 2016; Hoy et al. 2016; Kenny et al. 2016; Schwager et al. 2017; Gainett et al. 2021; Ontano et al. 2021). These advances have revolutionized modern views of chelicerate phylogeny and evolution, prompting reevaluations of historical paradigms of habitat transition and evolution of complex characters (Bond et al. 2014; Fernández et al. 2014; Sharma et al. 2014; Ballesteros and Sharma 2019; Ontano et al. 2021; Ballesteros et al. 2022).

Solifugae, variously known as “camel spiders” or “sun spiders”, is a mesodiverse group (in comparison with other arachnid groups) that currently includes ca. 1,200 species classified in 12 families and 144 genera (World Solifugae Catalog 2022). Notorious for their fearsome appearance, large chelicerae (relative to body size), and high bite force, (Meijden et al. 2012), solifuges are one of seven chelicerate orders that are commonly referred to as “the neglected cousins” in the arachnological community, due to a dearth of systematic and phylogenetic studies (Harvey 2002). Various characteristics of solifuges distinguish them from other arachnids, such as a robust pair of two-segmented chelicerae, adhesive structures on the termini of the pedipalps for prey capture, the presence of malleoli (“racquet organs”, used for chemoreception and analogous to the pectines of scorpions) on the ventral surface of leg IV (Brownell & Farley 1974; Cushing et al. 2005; Willemart et al. 2011), and an extensive tracheal system with additional stigmata (Franz-Guess & Starck 2016). The last of these is understood to facilitate the rapid running speed of solifuges, as well as their inhabitation of some of the driest habitats on Earth. Historically, solifuges were thought to be closely related to pseudoscorpions (Weygoldt and Paulus 1979; Shultz 2007), a hypothesis that has since been contradicted by sperm ultrastructure (Alberti and Peretti 2002), rare genomic changes and the ensuing placement of pseudoscorpions as the sister group of scorpions (Ontano et al. 2021); but see (Michalski et al. 2022). Solifugae has more recently been recovered as part of a clade with acariform mites (Poecilophysidea), and possibly also palpigrades (Cephalosomata) (Pepato et al. 2010; Ballesteros et al. 2022). A close relationship of these three orders is supported by the arrangement of the anterior body segments and the structure of the coxal glands, and is recovered by a subset of molecular analyses that have emphasized dense taxonomic sampling (Pepato et al. 2010; Dunlop et al. 2012; Ballesteros et al. 2019, 2022).

By contrast to their placement in higher-level chelicerate phylogeny, the internal relationships of Solifugae remain virtually unknown (Harvey 2002). A classification of the twelve extant families was proposed by Roewer (1934), based on overall similarity of characters that are, in turn, highly variable across genera and species; this classification largely remains in place (World Solifugae Catalog 2022). Beyond the incidence of poorly delimited higher-level taxa, a peculiarity of solifuge systematics is the concentration of diagnostic characters in the adult males of many lineages—in some groups, juveniles cannot be reliably assigned even to a specific family. As a result, few attempts have been made to infer phylogenetic relationships using either morphological or molecular datasets. Rhagodidae was suggested to be well-separated from the remaining solifuge families on the basis of its peculiar morphology, but this inference was not based on formal analyses and the polarity of these morphological characters is unknown (Roewer 1934). Investigations of specific character systems have documented a large diversity, such as the flagellum of the male chelicera and the histology of sperm ultrastructure, but these data have not been applied toward the goal of formal inference of evolutionary relationships (Klann et al. 2009, 2011; Bird et al. 2015). Although molecular data offer considerable advantages over anatomical datasets for phylogenetic study of enigmatic groups like Solifugae, few works have addressed internal relationships within this order. For example, RADseq data supported the monophyly of genus A recent analysis of *Eremocosta* and suggested post glacial colonizations in North American deserts (Santibáñez-López et al. 2021). A recent analysis of A recent analysis of Iranian solifuges based on one mitochondrial gene was able to sample seven families and the ensuing topology suggested the paraphyly of at least three families, but without significant nodal support for most interfamilial relationships (Maddahi et al. 2017). The only multilocus dataset applied to solifuge relationships examined the phylogeny of the North American family Eremobatidae and this four Sanger locus-based analysis uncovered extensive non-monophyly of constituent genera, together with limited nodal support for deep nodes (Cushing et al. 2015).

Given the presumably ancient diversification of Solifugae, as inferred from the appearance of crown group lineages by the Mesozoic and multiple fossil genera (Dunlop 2010, World Solifugae Catalog 2022), bridging the knowledge gaps in solifuge higher-level phylogeny requires the application of phylogenomic datasets and tissue sampling from rare, often aging, tissue collections. We therefore amassed a set of 120 solifuge species drawn from natural history collections around the world and sequenced these for a chelicerate-specific suite of ultraconserved elements (UCEs), an approach notable for its demonstrable capability to accommodate even highly degraded tissues. Here, we provide a robustly resolved phylogenomic tree of Solifugae, in tandem with molecular dating efforts to provide a temporal context to solifuge diversification.

## Results

We compiled a comprehensive data set with high-quality ultraconserved profiles spanning all extant chelicerate orders, represented by 129 taxa. In this data set, Solifugae was represented by 107 taxa, amounting to almost 10% of the global fauna (World Solifugae Catalog 2022), of which we generated ultraconserved element (UCE) libraries for 90 (84%) and *in silico* extracted UCEs from existing UCE libraries, transcriptomes or genomes for the remainder. Selection of outgroups prioritized the sampling of basal splits, to facilitate node calibration in molecular dating. For each of these thresholds for probe-to-library identity, we analyzed a family of matrices assembled by varying taxon occupancy thresholds, from 10% to 50% occupancy. In addition, to mitigate the impact of sparsely represented terminals, we created two additional families of matrices using only terminals with >100 UCE loci and varying taxon occupancy thresholds, both for the liberal and stringent thresholds for probe-to-library identity.

### Phylogenomic analyses

Maximum likelihood analyses based on 521 loci (25% occupancy; 65% probe-to-library identity threshold) recovered a tree topology that divided solifuge families into two major clades (which we refer as Clade I and II in the following text, see Figure 5). All families except Ammotrechidae and Daesiidae were recovered as monophyletic with support, barring Melanoblossidae, represented herein by a single terminal (Figure 2). To rule out the possibility of systematic bias driven by compositional heterogeneity, GC-base proportions were mapped on the 25% occupancy phylogeny. Only the outgroup terminal, the palpigrade *Eukoenenia spelaea* exhibited a high proportion of GC-content; however, excluding this taxon from the analysis did not affect the interfamilial relationships of Solifugae.

**Figure 1.**
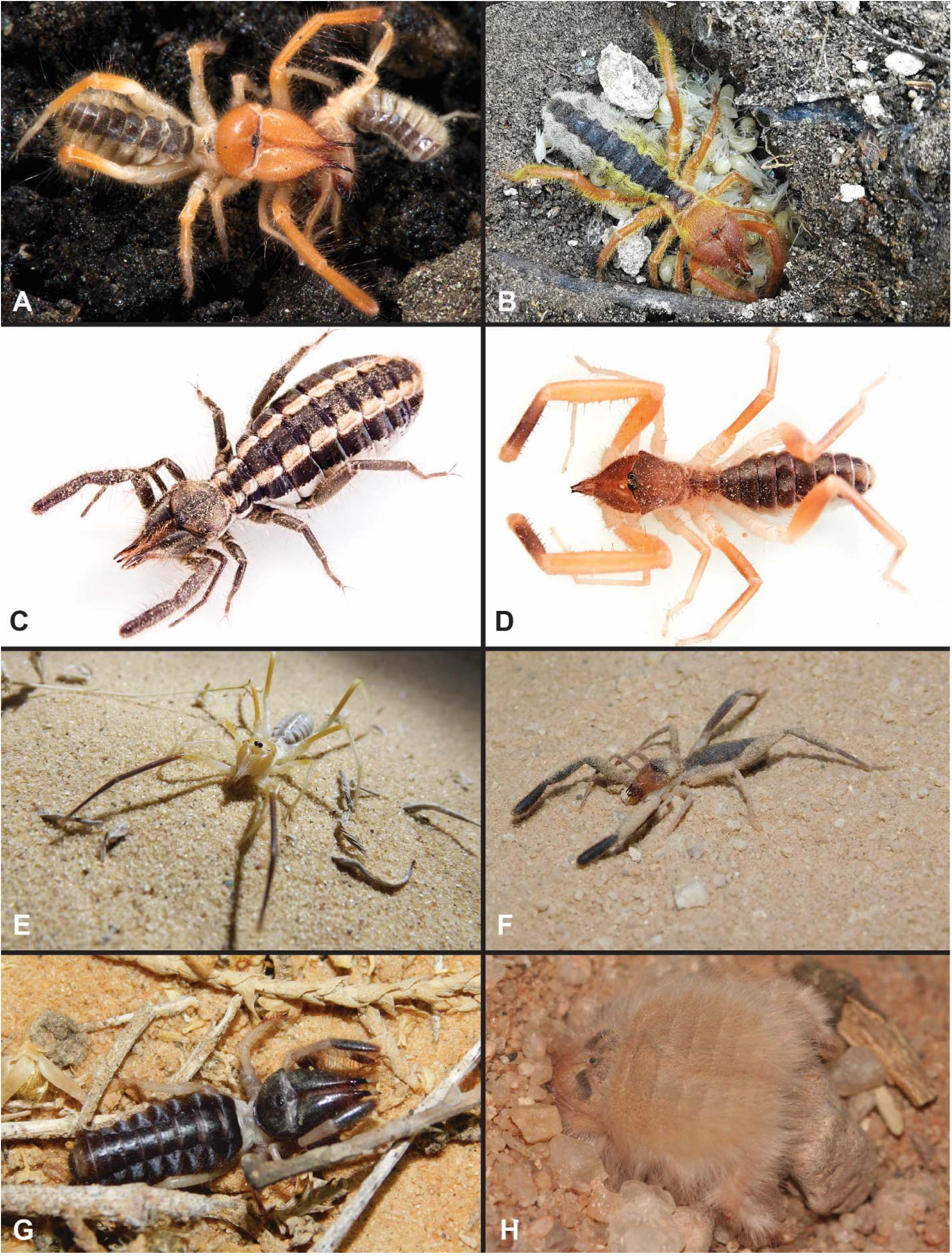
Live habitus of Solifugae. (A) Adult female of *Eremobates* (Eremobatidae) from Arizona, US. (B) Brooding female of *Galeodes* (Galeodidae) over a clutch of hatchlings, from Israel. (C) Adult female of *Mummucia* (Mummuciidae) from Chile. (D) Adult male of *Procleobis patagonicus* (Ammotrechidae) from Argentina. (E) Adult female of *Blossia* (Daesiidae) from Israel. (F) An unidentified Daesiidae from Israel. (G) An unidentified Rhagodidae from Israel. (H) An actively burrowing *Chelypus* (Hexisopodidae) from Namibia. Photographs: G. Giribet (A); I. Armiach (B, E, F, G); H. Iuri and A. Ojanguren-Affilastro (C, D); S. Aharon (H).

**Figure 2.**
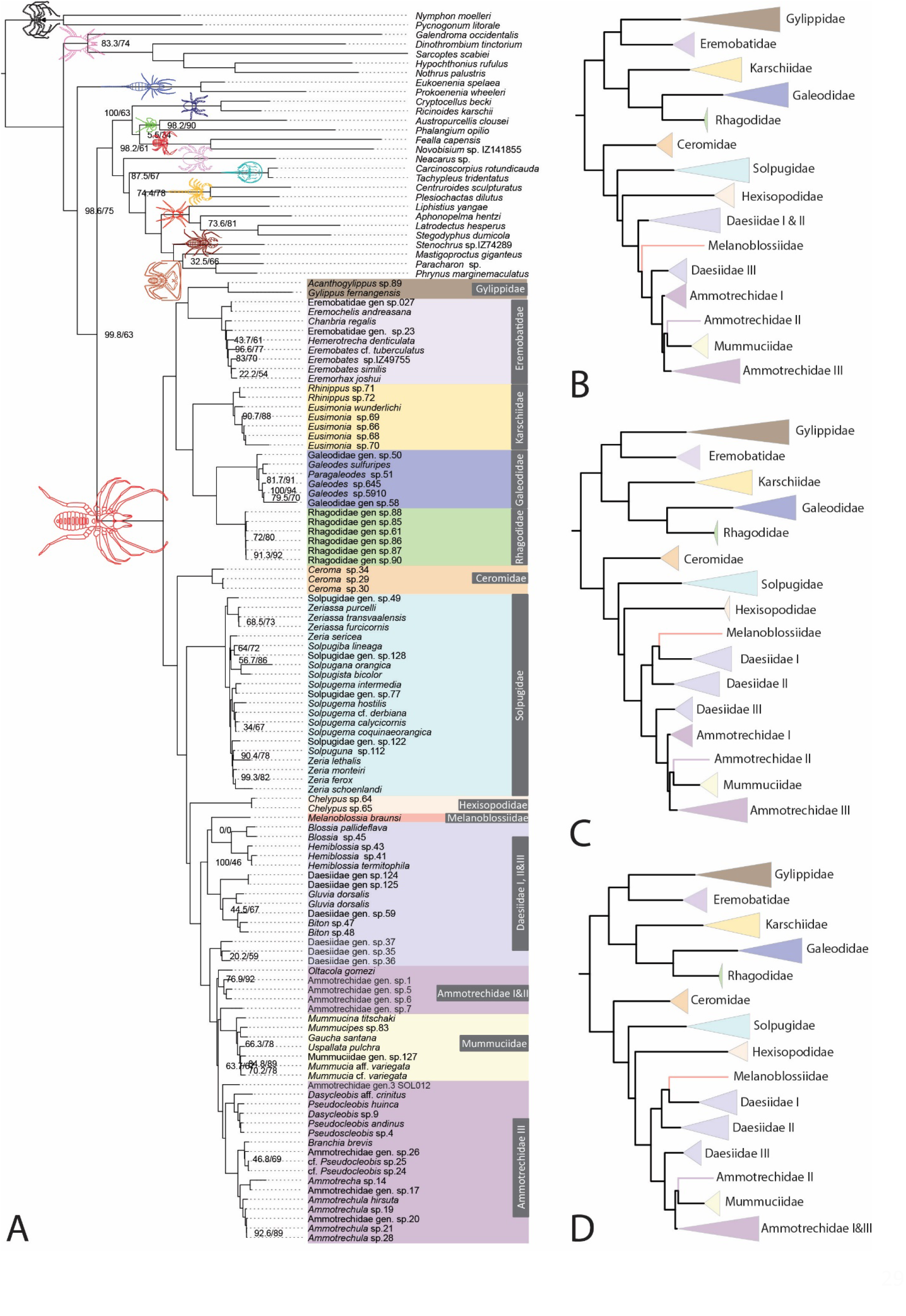
A maximum likelihood phylogeny of Chelicerata with all Solifugae family representatives using the 25% occupancy data set of ultraconserved elements.

The backbone tree topology of Solifugae was well-resolved and the basal split between the two clades was invariably recovered across analyses (Figures 2, 3). Mummuciidae was almost always nested within Ammotrechidae, rendering the family paraphyletic; only with a matrix filtered for taxon occupancy over 40% was Ammotrechidae recovered as monophyletic, albeit without support (Figures 2, 3). Melanoblossiidae was nested within the Daesiidae I clade with 25% and 35% occupancies (both poorly supported) and with 40% occupancy, the same relationship received high support (99% ultrafast bootstrap) whereas with 10% occupancy, it was placed as a sister group of a clade that included Daesiidae, Mummuciidae and Ammotrechidae (poorly supported).

**Figure 3.**
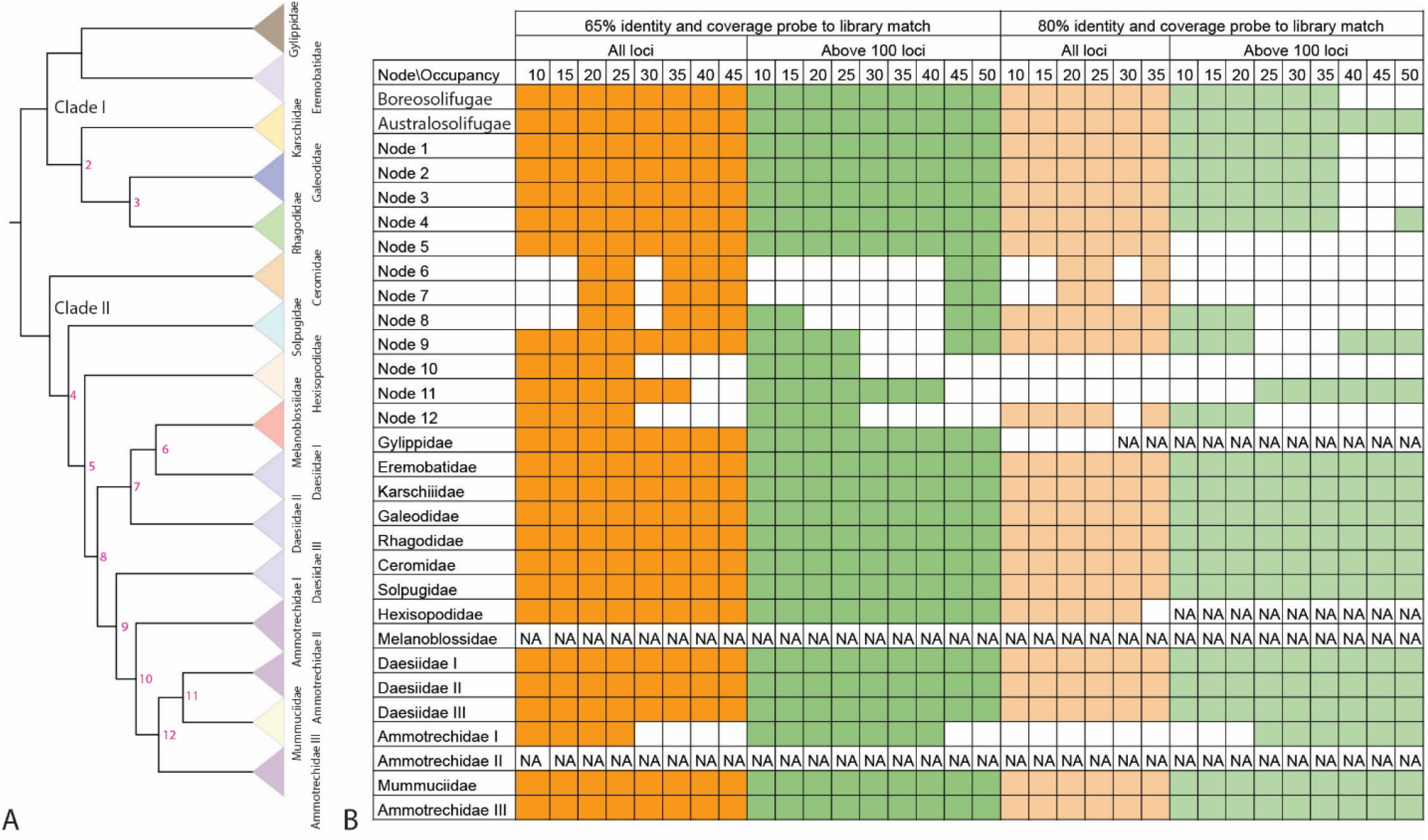
A comparison of interfamilial phylogenetic relationships of Solifugae compared across results of all analytical treatments. A. A maximum likelihood based phylogeny using the 25% occupancy data set (same as figure 2). B. Corroboration of relationships across different data sets at each node is shown in color palettes, alternative relationships are shown in white and NA indicates that either the family represented a single taxon or was not included in the analysis.

Across the >100 locus data sets, Mummuciidae was nested within Ammotrechidae in phylogenies inferred from 10-25% occupancy matrices. The remaining, more complete, but compact data sets (35-45%), recovered Mummuciidae as a sister group of Ammotrechidae with the exception of Ammotrechidae sp. SOL007. Only in the 50% occupancy data set was Ammotrechidae recovered as monophyletic (Figures S4-S9).

### Influence of stringent homology filtering

Matrices based on stringent filtering (80% identity and coverage probe-to-library match) discarded many loci, resulting in a more compact and denser data set across different occupancies. At occupancies above 20%, some terminals were dropped because the stringency of this filter did not recover any UCE locus for those taxa. Nevertheless, at least one taxon representing each currently recognized family was present in each occupancy data set, thereby not compromising the higher-level taxon sampling. The tree topologies recovered for matrices constructed under this threshold recovered similar relationships as previously reported. Gylippidae was recovered as paraphyletic in 10-25% occupancy matrices, but this is likely attributable to low locus representation (12 UCE loci for *Gylippus fernangensis*) at these occupancies. Among Solifugae, most interfamilial relationships were similar to that of the 25% occupancy phylogeny (Figure 3, S24-28). In the >100 locus data sets compiled using 80% homology mapping filter, most interfamilial relationships were similar to the 25% occupancy phylogeny. However, in ≥35% occupancy data sets, some outgroup taxa represented by very few loci were placed inside Solifugae. If these erroneous branches were to be pruned, solifuge interfamilial relationships continued to be robust even at 50% occupancy, which corresponds to a matrix of just 76 UCE loci.

### Divergence dating and biogeographic analyses

Divergence date optimizations calibrated with fossil age estimates in a least-squares (LSD2) framework, with and without ingroup fossils, recovered some non-overlapping age ranges. Diversification of the crown group Solifugae was estimated within the Carboniferous (344-305 Ma). Clade I diversified during the Permian-middle Triassic period (284-232 Ma), whereas Clade II diversified during the late Permian-late Triassic period (268-205 Ma). The MRCA (most recent common ancestor) of all families in Clade I, along with Ceromidae, Daesiidae III and Mummuciidae, originated in the Paleogene or earlier. The MRCAs of Melanoblossidae, Daesiidae I, Daesiidae II, Ammotrechidae I, Ammotrechidae II and Ammotrechidae III had a Cretaceous origin (Figure 4). Exclusion of the two Solifugae fossils recovered a more recent age range for all nodes, including a wider range of Carboniferous to Permian (334-250 Ma) age for the MRCA of Solifugae (Figure 4).

**Figure 4.**
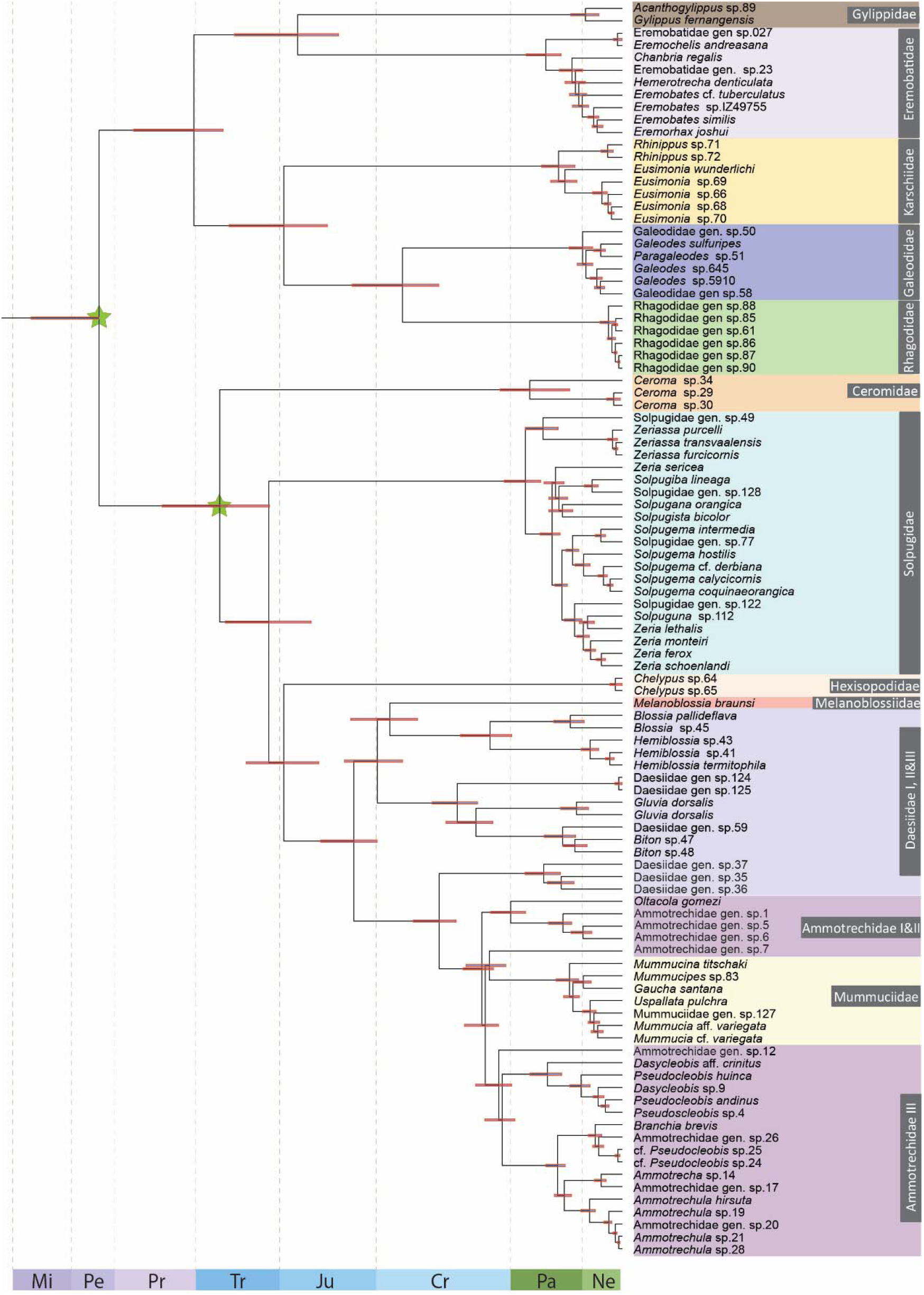
Fossil-calibrated dated phylogeny of Solifugae. Stars indicate fossil placements used for calibrating node ages. Outgroups removed for clarity.

To infer ancestral areas and reconstruct the biogeographic history of Solifugae, we analyzed the dated tree topology in tandem with five broad geographic areas using the RASP software (Figure 5). Given recent debates about the statistical comparability of models with and without the jump (*j*) parameter (Ree and Sanmartín 2018, Matze 2022), we trialed best-fitting models both with and without *j*. Historical biogeographic area optimizations were assessed by the best fitting model using RASP on our 25% occupancy data set. The model with highest AICc weight was the DEC+j (0.51), closely followed by DIVA-like+*j* (0.48). Among models without the jump parameter, the DEC model was closely followed by the DIVA-like model. Both DEC and DEC+*j* models recovered a combination of Turanian, African tropics, Neotropical and Mediterranean regions as the area for the most recent common ancestor (MRCA) of Solifugae (Figure 5). The ancestral area for Clade I was a combination of Turanian, Neotropical and Mediterranean (i.e., fragments of Laurasia), whereas for Clade II, the Afrotropics (i.e., a constituent of Gondwana) was recovered using both models (Figure 5). Most family distributions were limited to a single biogeographic region. One group of the polyphyletic Daesiidae (Daesiidae III) is distributed within Central Chile and the Patagonian (CCP) region and the other two groups have an African origin (Figure 5). At the crown node of the CCPian and Neotropical Ammotrechidae III clade, the DEC+*j* model optimized CCP as the ancestral area, which implied a dispersal event from CCP to the Neotropics. The exclusion of the jump parameter optimized a combination of CCP and Neotropical region as the ancestral area, thus implying a vicariance event dividing the descendant lineages (Figure 5). Seven dispersal events were implied by the DEC+*j* model (marked with red lines in Figure 5). These dispersals included three events between the Mediterranean region and African tropics; two between Neotropical and the CCP regions during the Paleogene period; one dispersal event from CCP to the Nearctic region during the Paleogene-Cretaceous boundary; and one African tropics to CCP regions during the Jurassic period were implied by the DEC+*j* model. However, the DEC model implied a vicariance event at these nodes. Overall, the DEC+*j* model favored a single area as origin where the descendent lineages have different regions implying a dispersal event. The exclusion of the jump parameter optimized a combination of two or more regions at the same ancestral nodes. Alternative coding of areas by continents for extant taxa recovered Asia and Africa as the ancestral areas for Clade I and Clade II, respectively. A further broader area coding by supercontinent recovered Laurasia as the ancestral area for Clade I and Gondwana for Clade II (Figures S30-31).

**Figure 5.**
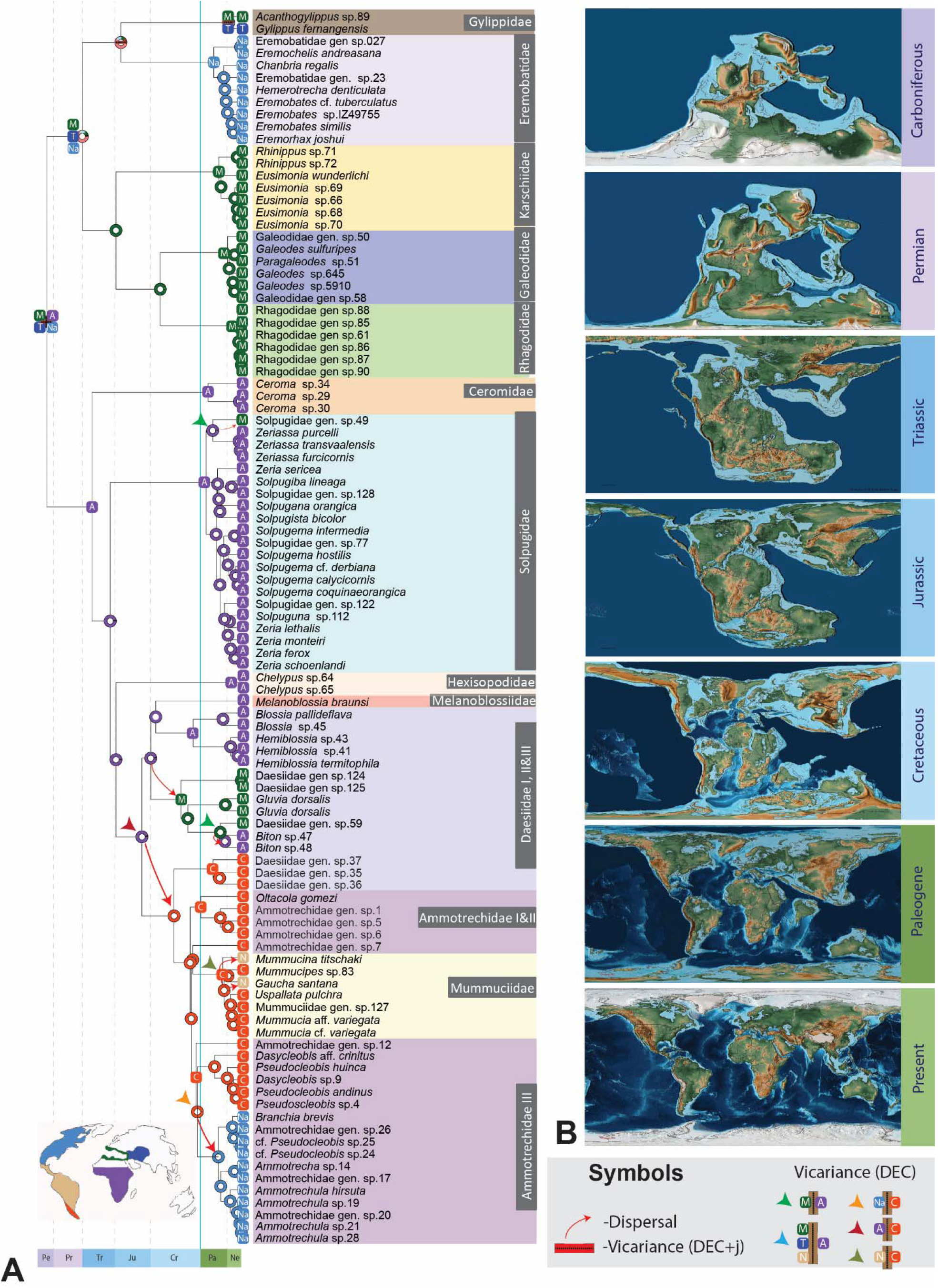
A. Biogeographic hypothesis obtained from the Dispersal-Extinction-Cladogenesis + jump parameter model of RASP analyses on the fossil-calibrated LSD2 analysis using the 25% occupancy data set. The dotted line in red indicates the Cretaceous Triassic boundary. B. Graphics for Pangean breakup from Scotese, C.R., 2016. PALEOMAP Project, http://www.earthbyte.org/paleomap--_paleoatlas--_for--_gplates/. Ancestral areas optimized for each family and higher-level nodes are marked at the respective MRCA nodes. Red arrows indicate dispersal implied by the DEC+j model whereas the colored arrowheads indicate alternative vicariance events implied by the exclusion of the *jump* parameter.

## Discussion

In a recent exploration of long branch attraction effects in chelicerate phylogeny, (Ontano et al. 2021) showed that taxonomic undersampling, and specifically, omitting the representation of basal nodes, exacerbated topological instability of fast-evolving arachnid orders like pseudoscorpions. Using a rare genomic change as a benchmark for phylogenetic accuracy, these analyses demonstrated that taxonomic sampling (especially of slowly-evolving and basally branching groups) outperformed other approaches to mitigating long branch attraction, such as increasing matrix completeness, using coalescent-aware species tree approaches, filtering by evolutionary rate, and implementing site heterogeneous models in concatenated matrices. Ontano et al. (2021) reasoned that applying this remedy to other fast-evolving orders may greatly stabilize chelicerate phylogeny, given that at least four orders exhibit a propensity for long branch attraction artifacts in Chelicerata (Ballesteros and Sharma 2019; Ballesteros et al. 2019; Ontano et al. 2022). This proposed remedy for long branch attraction is greatly potentiated by the availability of higher-level phylogenies for unstable groups, such as Acariformes (Klimov et al. 2017) and Palpigradi (Giribet et al. 2014). However, representation of solifuges in phylogenomic works has historically been driven entirely by the availability of sequence-grade tissues, rather than by phylogenetic representation, given that the internal phylogenetic structure of this order has never been fathomed (Borner et al. 2014; Sharma et al. 2014; Ballesteros and Sharma 2019; Arribas et al. 2020; Ballesteros et al. 2022; Ban et al. 2022). It is possible that this oversight may have underlain their known predilection for topological instability in some phylogenomic datasets (Sharma et al. 2014; Ballesteros and Sharma 2019; Ballesteros et al. 2019).

To redress this basic gap in our understanding of Solifugae, we generated the first higher-level phylogeny of the group, leveraging natural history collections worldwide to sample all extant families and reconcile a dated phylogeny against the fossil record. We obtained a robust backbone tree topology, with nearly all interfamilial nodes well-supported and recovered across an array of occupancy thresholds with support. Our results show that most solifuge families represent cohesive groups, with many families exhibiting high fidelity to specific biogeographic terranes. Curiously, we recovered the North American family Eremobatidae as the sister group of Gylippidae, which is restricted to the Old World (Central Asia and South Africa), with these two taxa in turn sister group to a large clade of Old World families (Galeodidae, Karschiidae, and Rhagodidae). This result was unanticipated because gylippids were previously considered part of Karschiidae (Roewer 1933), and because a North American sister group of this family implies a markedly disjunct distribution. However, this relationship is consistent with shared traits of cheliceral architecture and dentition in these two families (Bird et al. 2015).

Paralleling this outcome, the other groups of New World solifuges were also recovered as nested within a clade of Old World taxa (Ceromidae, Solpugidae, Hexisopodidae, and Old World daesiids). Our analyses consistently recovered a clade formed by the South American family Mummuciidae, the New World family Ammotrechidae, and the South American subset of the paraphyletic group Daesiidae (the “Daesiidae III” clade). Within each, we additionally found close relationships between geographically proximate subtaxa. These results suggest a prominent phylogenetic signature in solifuge biogeographic distributions, and across varying depths of the phylogeny.

Three relationships exhibited instability across analyses and invite further scrutiny. First, the fossorial group Hexisopodidae (commonly, “mole solifuges”), from southern Africa, exhibited some topological instability being recovered as either the sister group of Solpugidae in a minority of analyses, or with a clade formed by Daesiidae + Mummuciidae + Ammotrechidae. Both placements are partially supported by available morphological data. Specifically, analyses of solifuge sperm ultrastructure previously showed that hexisopodids and solpugids share similarities in the fine structure of spermatozoa, such as a conical acrosomal vacuole that is located within the chromatin body (Klann et al. 2011); an older work had also suggested that hexisopodids may constitute a derived subtaxon of Solpugidae (Hewitt 1919). But hexisopodids also exhibit digitiform protuberances of membranes and putative deposits of stored glycogen, which are similarly observed in Daesiidae and Ammotrechidae, in addition to Solpugidae (Klann et al. 2011). These data strongly accord with the recovery of these four families in a clade across our analyses, but we add the caveat that morphological data for solifuge sperm ultrastructure remain fragmentary; characteristics of the spermatozoa have been surveyed only in seven of the 12 families to date (Klann et al. 2009, 2011). In particular, data on the sperm ultrastructure of Mummuciidae, a member of this group, are not presently available.

The second relationship that exhibited instability was the placement of the sole exemplar of Melanoblossidae; a family with a disjunct distribution that encompasses southern Africa and Southeast Asia. A minority of analyses recovered *Melanoblossia* as the sister group of the Old World Daesiidae, whereas the majority recovered this terminal as derived within the Old World daesiids. As Melanoblossidae was previously considered a subfamily of Daesiidae (Roewer 1933), we cannot rule out a nested placement of this taxon. As with the hexisopodids, this topological instability across analyses stems from limitations in locus representation for *Melanoblossia*, which confers a high proportion of missing data for these terminals in the phylogenetic matrices we analyzed. Future investigations of hexisopodid and melanoblossid placement must emphasize deeper sampling and sequence coverage for these two taxa.

A final relationship that exhibited instability across analyses was the placement of Mummuciidae. While mummuciids were typically recovered as nested within Ammotrechidae, we did recover this clade as the sister group of the ammotrechids in a minority of analyses (albeit without support). As with the previously discussed case, mummuciids were previously a subtaxon of Ammotrechidae, and thus a nested placement of this putative family would not be unprecedented (Roewer, 1933). Given the complexity of ammotrechid subfamilies (a subset of which were sampled here), we submit that future systematic revisions of the South American solifuges must consider mummuciids as potentially derived members of Ammotrechidae.

With regard to biogeography, the chronological sequence of Pangean fragmentation is well-documented, and numerous cases of biotic distributions and divergence time estimates have been shown to retain the signature of supercontinental breakup (e.g., Sanmartín and Ronquist 2004; Giribet et al. 2012; Mao et al. 2012; Murienne et al. 2013b). Our biogeographic analysis revealed the general pattern that the distribution of most solifuge clades is delimited by single biogeographic regions. Ancestral area optimizations in the internal nodes and the distribution of extant species (e.g., near-absence on oceanic and Darwinian islands) suggest that solifuges are habitat specialists adapted to specific environments, being largely confined to xeric habitats.

Like many arachnid orders, the crown group of Solifugae dates to the Carboniferous or earlier, as reflected by its fragmentary fossil record (World Solifugae Catalog 2022; Dunlop 2010). Most solifuge families and genera are relatively young, diversifying soon after the Cretaceous-Paleogene extinction event, particularly in Clade I. While this may reflect an artifact of limited sampling, one possibility for the observation of relatively young ages of family-level clades is that the recent aridification and the opening of comparatively young deserts may have opened new ecological niches for Solifugae, driving the diversification of this arid-adapted arachnid group by the mid-Cenozoic (Zhang et al. 2014; Zheng et al. 2015; Cushing et al. 2015). Their absence on dry continental landmasses such as Australia may be attributable to the recent timing of aridification of the Sahul Shelf, as well as the mesic and cooler past conditions of the Australian interior (Byrne et al. 2011). However, assessing this hypothesis using parametric tests requires extensive sampling of extant diversity, in tandem with consideration of variance in the ages of deserts worldwide. Targeted approaches like UCE sequencing offer the promise of renewed utility of natural history collections and revitalized prospects for hypothesis testing with poorly studied taxa.

We emphasize that various subtaxa were not included in this study, which is aimed specifically at higher-level relationships between the families. Given the fidelity we observed between solifuges and the terranes that they inhabit, we anticipate that future sampling efforts will identify additional cases of non-monophyletic taxa from regions not represented in this study (e.g., the Indian subcontinent; eastern China), which will require intensive rounds of systematic revision under a phylogenomics lens. Such genomics-driven revisionary efforts have demonstrated marked success and efficiency in modernizing the classification and evolutionary analysis of comparably diverse groups, such as scorpions and harvestmen (Sharma et al. 2015; Derkarabetian et al. 2018; Benavides et al. 2019b; Santibáñez-López et al. 2019, 2020).

## Conclusions

The phylogenetic tree topopology presented herein bookends a ca. 20-year gap of available higher-level phylogenies for the orders of Chelicerata (Harvey 2002). The relationships inferred herein are anticipated to aid taxonomic revisionary efforts, and revitalize morphological comparative studies and biogeographic efforts within this understudied group. Future investigations of solifuge evolutionary history must emphasize the role of global perturbations of the past and the downstream effects of climatic cycles on diversification and range expansion of these cryptic, long-neglected arthropods.

## Materials and Methods

### Species sampling

Specimens sequenced for this study were collected from field sites as part of our recent collecting campaigns, as well as contributed by collections of the Museo Argentino de Ciencias Naturales “Bernardino Rivadavia”, Buenos Aires, Argentina (MACN); The National Natural History Collections, The Hebrew University, Jerusalem, Israel (NNHC); the Denver Museum of Nature & Science, Colorado, United States (DMNS); the National Collection of Arachnida, Agricultural Research Council, South Africa (ARC-PCP), and the Zoological Museum Hamburg (ZMH). Collecting permits for study taxa were issues to different laboratories over several years; permitting data are available upon request from the authors. For *de novo* sequencing of UCEs, sampled exemplars of each of the 12 described extant families, as follows: 18 Ammotrechidae, six Ceromidae, 15 Daesiidae, five Eremobatidae, nine Galeodidae, two Gylippidae, four Hexisopodidae, eight Karschiidae, five Melanoblossidae, six Mummuciidae, ten Rhagodidae, and 25 Solpugidae. To this dataset, we added UCE loci from five published solifuge transcriptomes, comprising one Ammotrechidae, two Eremobatidae and two Galeodidae

Outgroup sampling was influenced by previous works that have inferred various possible placements of Solifugae in the chelicerate tree of life, such as a sister group to Acariformes and Palpigradi, or part of a clade with Riniculei, Opiliones, and Xiphosura (Pepato et al. 2010; Sharma et al. 2014; Ballesteros and Sharma 2019; Ballesteros et al. 2019, 2022). Given this instability across studies, we sampled 2-3 terminals of every extant chelicerate order, prioritizing the sampling of basal splits in each outgroup order. Outgroup datasets were drawn from published transcriptomes and from our previous UCE assemblies that were captured with the same probe set. All tree topologies were rooted with Pycnogonida.

### Ultraconserved element sequencing and phylogenomic analyses

For newly sequenced specimens, 1-4 legs or tissue teased from a single chelicera from single specimens were used for DNA extractions using the DNeasy™ Blood and Tissue kit and the QIAamp DNA Mini kit (Qiagen Inc., Valencia, CA). Libraries were prepared and enriched following protocols outlined by (Kulkarni et al. 2020) and Miranda et al. (2022). All pools were enriched with the Spider2Kv1 probe set (Kulkarni et al. 2020) following the myBaits protocol 4.01 (Arbor Biosciences). Sequencing was performed on an Illumina NovoSeq platform. Assembly, alignment, trimming and concatenation of data were performed using the PHYLUCE pipeline (publicly available at https://phyluce.readthedocs.io/en/latest/). UCE contigs were extracted using the Spider2Kv1 probe set (Kulkarni et al. 2020) to target 2,021 UCE loci. To augment the UCE dataset with RNASeq datasets, we followed the assembly, sanitation, reading frame detection, and UCE retrieval pipeline outlined by Kulkarni et al. (2021). Homology screening was performed by employing liberal (65%) and stringent (80%) probe-to-library identity and coverage mapping thresholds as suggested by Bossert & Danforth (2018).

### Phylogenomic analyses

The assembly, alignment, trimming and concatenation of data were done using the PHYLUCE pipeline (publicly available at https://phyluce.readthedocs.io/en/latest/). We assessed the sensitivity to data completeness by applying gene occupancy with successive increments of 5% (10% onwards) to where the locus count dropped below 50 UCE loci.

Orthologous and duplicate loci were screened with the minimum identity and coverage of 65%. Phylogenetic analyses were performed on the unpartitioned nucleotide data using IQ-TREE (Nguyen et al. 2015) version 2. Model selection was allowed for each unpartitioned dataset using the TEST function (Kalyaanamoorthy et al. 2018, Hoang et al. 2018). Nodal support was estimated via 2,000 ultrafast bootstrap replicates (Hoang et al. 2018) with 10,000 iterations. To reduce the risk of overestimating branch support with ultrafast bootstrap due to model violations, we appended the command -bnni. With this command, the ultrafast bootstrap optimizes each bootstrap tree using a hill-climbing nearest neighbor interchange (NNI) search based on the corresponding bootstrap alignment (Hoang et al. 2018).

### GC-content

GC content can influence the phylogenetic relationships reconstructed using genome scale data (Benjamini and Speed 2012). To address this, we computed GC content for each taxon in the concatenated alignment using a custom script paired with BBMap (https://github.com/BioInfoTools/BBMap).

### Phylogenomic dating

As no molecular phylogeny of the order exists and the solifuge fossil record is sparse, the timeframe of solifuge diversification is effectively unknown. To provide a temporal context to the divergences we inferred, we performed divergence time estimation using a least-squares approach, LSD2 method (To et al. 2016) which uses a least-squares approach based on a Gaussian model and is robust to uncorrelated violations of the molecular clock. LSD2 requires at least one fixed date, so we used an absolute calibration of 314.6 Ma for the crown Orthosterni fossil, *Compsoscorpius buthiformis*. We optimized the fossil information-based calibrations on the tree topology inferred from the 25% occupancy data set for the basis for divergence time estimation, implementing uniform node age priors to accommodate the scarcity of terrestrial chelicerate fossils. The root age was set to a range of 550–600 Mya, following Wolfe et al. (2016). The crown age of Solifugae was constrained using a minimum age bound of 305 Ma, based on the Carboniferous fossil *Prosolpuga carbonaria* (Petrunkevitch, 1913). The stem age of Ceromidae was constrained using a minimum age bound of 115 Ma, based on the Cretaceous fossil *Cratosolpuga wunderlichi* (Selden and Shear 2016). The recently described Burmese amber fossil *Cushingia ellenbergeri* was not included for calibration, characters of this species potentially accord with a melanoblossid, a gylippid, or a rhagodid identity, which precludes its use for calibrating a specific node (Dunlop et al. 2015; Bartel et al. 2016). Outgroup nodes were calibrated using the oldest unambiguous fossils representing those clades.

### Ancestral area reconstruction

We reconstructed ancestral areas on internal nodes of the dated preferred tree and the combined tree using the package BioGeoBEARS (Matze, 2013) implemented in RASP 4.0 (Yu et al., 2020). Each of the terminals was assigned to one of the following biogeographic regions: Turanian, African tropics, Neotropical, Nearctic, Central Chile-Patagonia and Mediterranean. We chose this scheme of area coding based on the distribution of the extant solifuges and following commonly used area definitions from the literature. Some taxa such as ammotrechids and eremobatids are distributed in the United States and Mexico. For these taxa, we coded the area as Nearctic following the delimitation of this region by Escalante et al. (2010). Additionally, to assess the influence of area coding, we alternatively coded areas by continents (corresponding to their geological plate); and more coarsely, as either Laurasia or Gondwana, based on the geological origin of those areas.

## Acknowledgements

We are grateful to Nuria Macias (Departamento de Zoología de la Universidad de La Laguna) for providing a specimen of *Eusimonia wunderlichi* from the Canary Islands, and to Marc Domènech and Miquel Arnedo (University of Barcelona) for providing a specimen of *Gluvia dorsalis* from Spain. Photographs of live specimens were kindly provided by Igor Armiach Steinpress and Shlomi Aharon. Sequencing was performed at the Biotechnology Center of the University of Wisconsin-Madison.

